# Adaptive Optical Coherent Raman Imaging of Axons through Mouse Cranial Bone

**DOI:** 10.1101/2022.09.14.507912

**Authors:** Jong Min Lim, Seokchan Yoon, Seho Kim, Youngjin Choi, Jin Hee Hong, Wonshik Choi, Minhaeng Cho

**Author notes:** Both authors contributed equally to the manuscript (Jong Min Lim, Seokchan Yoon).

## Abstract

Coherent Raman scattering imaging has provided inherent chemical information of biomolecules without the need for any external labels.^1–3^ However, its working depth in deep-tissue imaging is extremely shallow because both the intrinsic scattering cross-section and image contrast are so small that even weak perturbation of the pump and Stokes beam focusing by the complex tissue causes the loss of the resolving power.^4,5^ Here, we propose a deep-tissue coherent Raman scattering (CRS) microscopy equipped with an advanced adaptive optics (AO) system measuring complex tissue aberration from elastic backscattering. Using this label-free AO-CRS microscopy, we demonstrate the vibrational imaging of lipid-rich substances such as myelin inside the mouse brain even through the thick and opaque cranial bones.

---

Recent progress of coherent Raman scattering (CRS) microscopy that utilises inherent chemical contrast without any exogenous labels promises wide applications to understand genuine characteristics of biomolecules in living organisms.^1–3^ Since the coherent Raman process provides more enhanced signal intensity than the spontaneous Raman technique, it is capable of acquiring consecutive images toward a real-time observation of bio-molecular dynamics.^6,7^ Quantitative and qualitative assessments of biomolecules such as lipid, glucose, protein, and DNA in living organisms have been demonstrated even in situ.^8^ In addition, it allows the investigation of biomolecules in depth as the nonlinear optical process is generated only at the tight focus.

However, the penetration depth of CRS microscopy is much shallower than other nonlinear optical imaging techniques such as two-photon and three-photon microscopy.^4,5^ The scattering cross-section of the CRS process is many orders of magnitude smaller than other nonlinear processes relying on the labeling, and its image contrast is lower due to the background signals from the bulk tissues. Therefore, even the weak perturbation of the tight focusing by the spatially heterogeneous bio-structures can substantially reduce the signal intensity. This has limited its applicability to observing surfaces of biosystems or relatively transparent cases such as the animal ear.^9^ In order to overcome the penetration depth limit of CRS microscopy, generally known as 20 – 100μm in various tissue samples, several approaches have been suggested so far.^10,11^ For alleviating the photon scattering effect, a tissue clearance method removing intracellular components and a frequency red-shift of the incident beams towards the NIR region were suggested.^12–14^ Also, there have been attempts to insert a miniature microscope objective close to the target biological system. More recently, a combination of various nonlinear processes, including CRS with a machine learning algorithm, has been used to increase the penetration depth in brain tissues.^15,16^

Adaptive optics (AO) could provide an ideal solution for deep-tissue CRS imaging because it is a pure optical solution causing neither modification nor damage to the sample. AO uses wavefront shaping devices such as a deformable mirror and a liquid-crystal spatial light modulator (SLM) to compensate for the aberration and recover a tight focus in deep tissue.^17,18^ It has been widely used for increasing the imaging depth of various nonlinear imaging modalities. AO microscopy can be classified by the way to find the sample-induced aberration. One approach is sensorless AO, which controls the wavefront shaping device in a way that optimises image quality metrics such as image sharpness and brightness.^19–21^ With its simplicity of implementation, sensorless AO has been employed in CRS microscopy to compensate for sample- and system-induced aberrations.^22,23^ Still, this AO technique has a limitation that aberration should be mild enough for the sample to be initially visible for evaluating quality metrics, meaning that the imaging depth increase is marginal. The other approach is wavefront sensing AO, which directly measures the wavefront of returning waves to determine the sample-induced aberrations.^24–27^ For example, a Shack-Hartmann wavefront sensor is used to measure the wavefront of the fluorescence signal in multi-photon imaging. However, this approach is not suitable for CRS imaging because the signal level is too low to measure its wavefront. We noted that the wavefront sensing AO based on the measurement of the intrinsic elastic backscattering could be well suited for deep-tissue CRS imaging.^28–31^ Since the wavefront sensing process is conducted without any labels, it preserves the label-free imaging capability of the CRS imaging. Furthermore, it can be applicable at depths CRS imaging is entirely disrupted as long as the working depth of wavefront sensing is deeper than that of CRS imaging.

Here, we employ the closed-loop accumulation of single scattering (CLASS) system,^31–33^ one of the most advanced wavefront sensing AO systems, to the epi-detection coherent Raman scattering microscopy for vibrational imaging of biomolecules deep within complex tissues. CLASS measures a time-gated reflection matrix of the intrinsic elastic backscattering from the tissue to identify complex sample-induced aberrations. Its working depth is sufficiently high enough to quantify tissue aberrations even through the intact skull.^33^ In the backbone of the CRS microscopy system, a unique common-path off-axis digital holographic system is added for the robust measurement of the reflection matrix. The identified aberration map by the reflection matrix is transferred to the SLM in the excitation beam path of the CRS microscopy to compensate for the tissue aberrations of both the pump and Stokes beams. For the optimal aberration correction, the aberration of the pump beam responsible more for the CRS signal generation than the Stokes beam is measured. We conduct coherent anti-Stokes Raman scattering (CARS) imaging of the brain cortex through a thinned skull and visualize many myelinated axons initially invisible with a spatial resolution of about 300 nm, close to the diffraction-limited resolution accounting for the nonlinear excitation. There is a ten-fold enhancement of the pump point spread function (PSF) Strehl ratio, and the achieved image contrast of myelin segments beneath the cranial bone is comparable to that taken for the bare tissue.

The schematic of the experimental setup is shown in Fig. 1 (see the detailed layout in Fig. S1), which is composed of CRS and CLASS sections. In the CRS section, pump and Stokes pulses are overlapped in space and time. A half waveplate is inserted to generate *p*- and *s*-polarized pulses, which are used for sample and reference beams, respectively, in the CLASS scheme. Both the pump and Stokes pulses travel through the SLM. Then, they are sent to the microscope and steered with a pair of Galvano mirrors for the 2D lateral scanning. The SLM is initially used as a flat mirror for aberration measurement, and an aberration correction map is displayed for the deep-tissue CRS imaging. The *s*-polarized pulses are reflected by a polarizing beam splitter (PBS1) toward the reference mirror (RM). They will be used as reference pulses in the CLASS system. The transmitted *p*-polarized pump and Stokes pulses are focused on the sample via a microscope objective (60x, 1.0 NA) to generate a CARS signal collected by a dichroic mirror (DCM2) and detected by PMT. In our system preparation, we make sure that the foci of the pump and Stokes pulses are matched within 1 μm precision (less than 2λ) using the knife-edge measurement (Fig. S2).

**Fig. 1:**
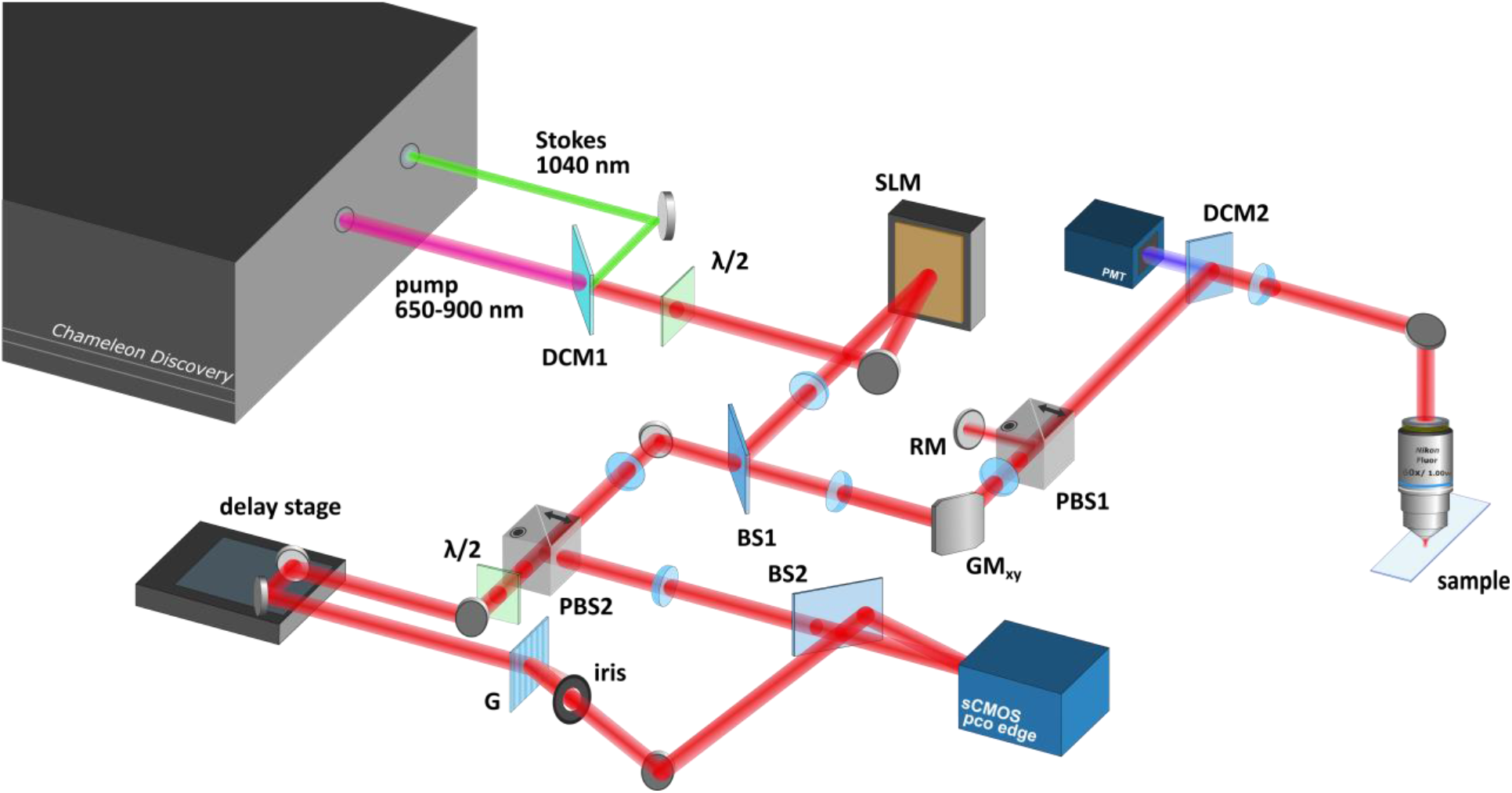
Schematic of the adaptive optical coherent Raman scattering (AO-CRS) microscopy. The setup is composed of two sections: CLASS and CRS. DCM1, 2: dichroic mirrors, λ/2: half-wave waveplate, SLM: spatial light modulator, BS1, 2: beam splitters, GM: Galvanometer scanning mirror, PBS1, 2: polarizing beam splitter, RM: reference mirror, PMT: photomultiplier tube, PD: photodiode, G: grating.

The CLASS section measures the tissue-induced aberrations by recording the electric field maps of the elastic backscattering of the pump pulse from the sample. To this end, we construct a unique common-path off-axis digital holographic system where sample and reference beams share Galvano scanning mirrors. This configuration eliminates the unwanted phase induced by the mismatch of the pivot in the scanning mirrors to ensure the recording of electric field maps solely from the sample. The elastic backscattering of the pump-Stoke pair from the sample and reference pulses reflected by RM are recombined at PBS1 and sent to the CLASS section through a beam splitter (BS1). The sample and reference beams are separated and recombined by PBS2 and BS2, respectively. The difference in their optical paths is compensated by the mechanical delay stage in the reference beam path. A diffraction grating is inserted in the reference beam path along with an iris diaphragm choosing the first-order diffracted beam to generate an off-axis interference image in the camera. The complex electric-field maps of the backscattered sample pulses are reconstructed from the recorded interference images by means of a Fourier-transform method.^33^

A set of complex field maps is recorded while scanning the focused beam at the sample to construct a reflection matrix, and the single-scattering correlation algorithm is applied to the matrix to identify the sample-induced aberration map.^20^ While one can measure the aberration maps of any of the pump and Stokes pulses, we choose to measure short-wavelength pump pulses considering that they experience more pronounced aberrations than the Stokes pulses, and the square of pump intensity is involved in the CARS signal generation. The aberration correction map is displayed on the SLM placed at a plane conjugate to the pupil plane of the objective lens to simultaneously compensate for the sample-induced aberrations experienced by pump and Stokes pulses. This beam wavefront control by the SLM enables us to generate a tight focus at a sample plane located deep within tissues. SRS imaging channel is also prepared with a photodiode detector measuring stimulated Raman loss (SRL) of pump intensity (Fig. S1).

We validate the performance of our AO-CRS microscope by investigating the pump and Stokes PSFs and CARS signal intensities depending on the degree of aberration correction. An artificial aberration layer is placed on top of the 1-μm polystyrene beads (Fig. 2a). The CARS images are measured by targeting CH stretching vibration of polystyrene (3050 cm^-1^) by setting the pump and Stokes wavelengths to 789 and 1040 nm, respectively. The upper and lower images in Fig. 2b show the CARS images before and after the aberration correction, respectively. The inset in the lower image shows the aberration correction map, *ϕ_c_*(*k_x_,k_y_*), displayed on the SLM. It is noteworthy that the polystyrene beads are completely invisible before the correction, and they are made visible with high contrast after the correction, confirming the detrimental effect of aberration and proving the validity of the aberration correction. Next, we control the degree of aberration correction by displaying *α* × *ϕ_c_*(*k_x_,k_y_*) on the SLM with 0 ≤ *α* ≤ 1 and measure the PSFs of the pump and Stokes pulses at the sample plane (Fig. 2c) using a transmission-mode detection (Fig. S1). For instance, *α* = 1 and *α* = 0 correspond to full correction and no correction, respectively. The first row in Fig. 2c shows the residual aberration map of the pump pulses measured by the CLASS system for various *α*’s. With the increase of *α*, both the pump (second row) and Stokes (third row) PSFs are getting narrower, and their peak intensities increase accordingly. The width of pump PSF at *α* = 1 is 400 nm, close to that of the diffraction-limited PSF (395 nm), supporting the precision of the aberration correction. The relative pump Strehl ratio defined by the peak PSF intensity with respect to that of *α* = 1 is indicated at the upper right of each figure panel, which shows that peak pump intensity is increased by a factor of 6.25 after full correction. The Stokes PSF is slightly broadened such that its width is 1.3 times that of the diffraction-limited PSF. A slight focus mismatch between the pump and Stokes beams due to the system dispersion causes a factor of 1.2 broadening (Fig. S2). The residual discrepancy arises because the aberration measurement and correction are designated for the pump pulse. Still, there is a substantial peak intensity enhancement (1.97 times) of peak Stokes intensity from *α* = 0 to *α* = 1, supporting the effectiveness of the single SLM correction for the pump and probe beams with two different wavelengths. Considering that the peak intensity of PSF is inversely proportional to the square of its width, the peak intensity of the Stokes beam would have been enhanced by a factor of 3.3 if one more SLM were used for independently correcting the Stokes beam. The CARS images of the polystyrene bead in the fourth row of Fig. 2c and their line profiles in Fig. 2d along the dashed lines show the steady increase of signal intensity in accordance with the increase of the pump and Stokes Strehl ratios.

**Fig. 2:**
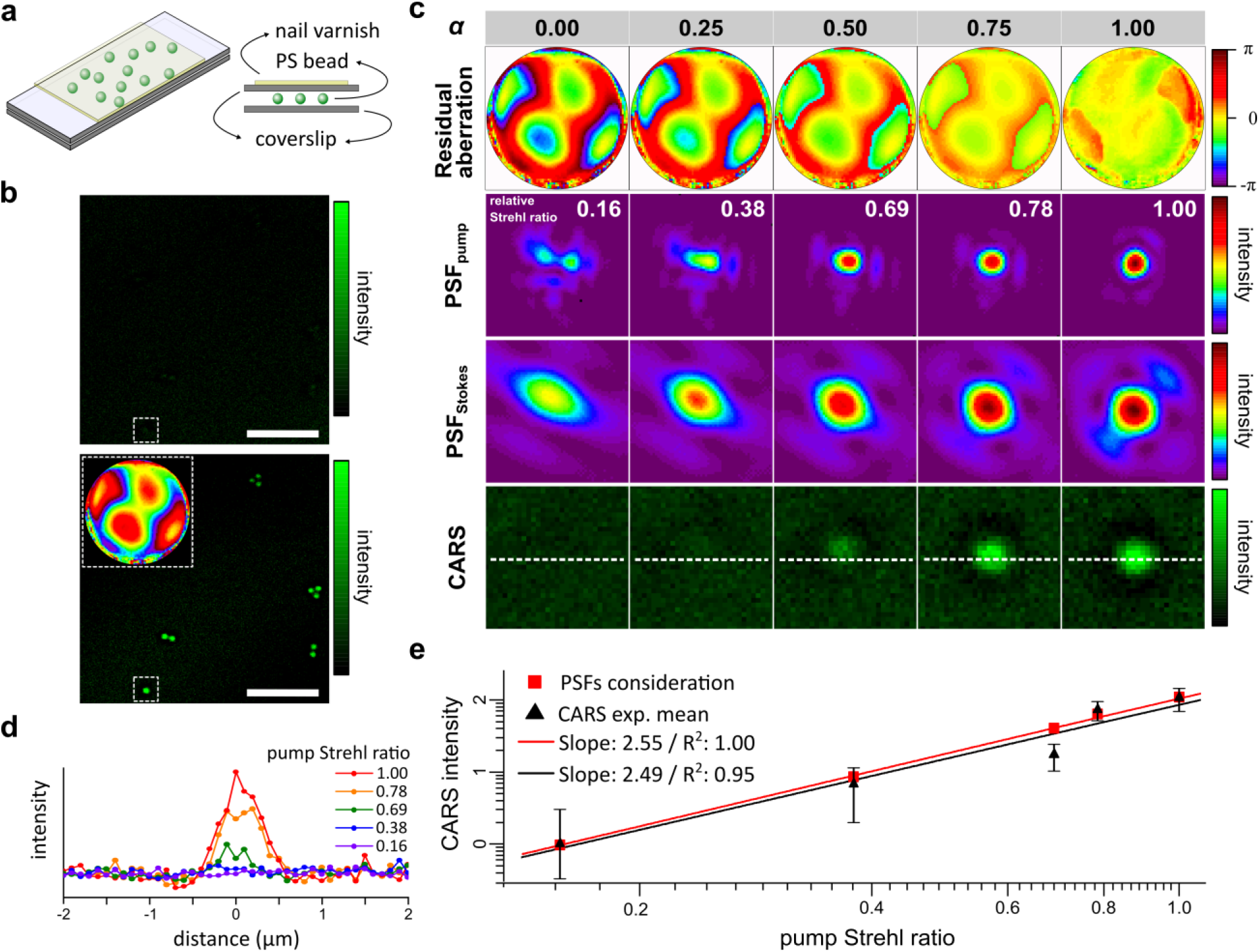
Experimental validation of the aberration correction capability. **a,** Schematic presentation of the artificial target. The 1-μm polystyrene beads were dispersed in between two coverslips (100 μm thickness), one of them placed on the top is coated with nail varnish for an artificial aberration. **b**, Representative CARS and CLASS-CARS images of polystyrene beads when the aberration was fully corrected. Inset in the lower image is the aberration correction map displayed on the SLM. **c**, Residual aberration maps (first row), resulting PSFs of the pump beam (second row), those of Stokes beams (third row), and the CARS images of a polystyrene bead (fourth row) depending on the degree of aberration correction. **d**, Line-cut profiles along the dashed lines of CARS images in (c) for different pump Strehl ratios. **e**, CARS intensity versus pump Strehl ratio obtained from (c) (triangular dots). CARS intensities estimated by the pump and Stokes PSFs (red square dots) were plotted as a point of reference. The scale bars in (c) indicate 10 μm. Aberration maps are the phase retardations in the pupil plane. Color bars in aberration maps: phase in radians.

We validate that CARS signal intensity is proportional to the square of the pump Strehl ratio multiplied by the Stokes Strehl ratio by polystyrene bead image, a point-like object. The peak intensity of the CARS signal is plotted as a function of the pump Strehl ratio (triangular dots in Fig. 2e). For comparison, the expected CARS signal intensities are estimated by the Strehl ratios of the pump and Stokes PSFs in the second and third rows, respectively, in Fig. 2e (red square dots). As expected, the slope of the CARS intensity (2.49) is close to that estimated by the pump and Stokes PSFs (2.55). From this observation, it is legitimate to state that the CARS signal is increased by a factor of 77 with the full aberration correction.

Point particles are rather ideal targets that seldom exist in realistic imaging conditions. Instead, we investigate the performance of our system for imaging a spatially extended target under severe aberration. A gold-deposit resolution target is covered with a PDMS polymer layer, and a crumpled vinyl film introducing the complex optical aberrations is placed on the top (Fig. 3a). The identified aberration map is applied to the reflection matrix to computationally correct the aberration of the time-gated confocal reflectance image reconstructed from the reflection matrix, which results in the enhanced signal intensity and contrast (Fig. 3b). The aberration map (Fig. 3c) is obtained from the time-gated reflection matrix recorded for the elastic backscattering of the pump pulses from the gold layer. We then obtained the CARS images by targeting the CH vibration of the PDMS polymer at 2905 cm^-1^. Before the correction, image sharpness is low, and fine structures in the central region are missing (Fig. 3d). Displaying the input aberration correction map to the SLM, we obtained the aberration-corrected CARS image (Fig. 3d and Fig. S3), which shows the enhanced contrast and resolution. The circular cut profiles of CARS images for identical spatial positions show an 8-fold enhancement of contrast, almost an order of magnitude improvement. Achieved spatial resolution after the correction is 0.7 μm, which is slightly lower than the diffraction-limited resolution of 0.4 μm (Fig. S4).

**Fig. 3:**
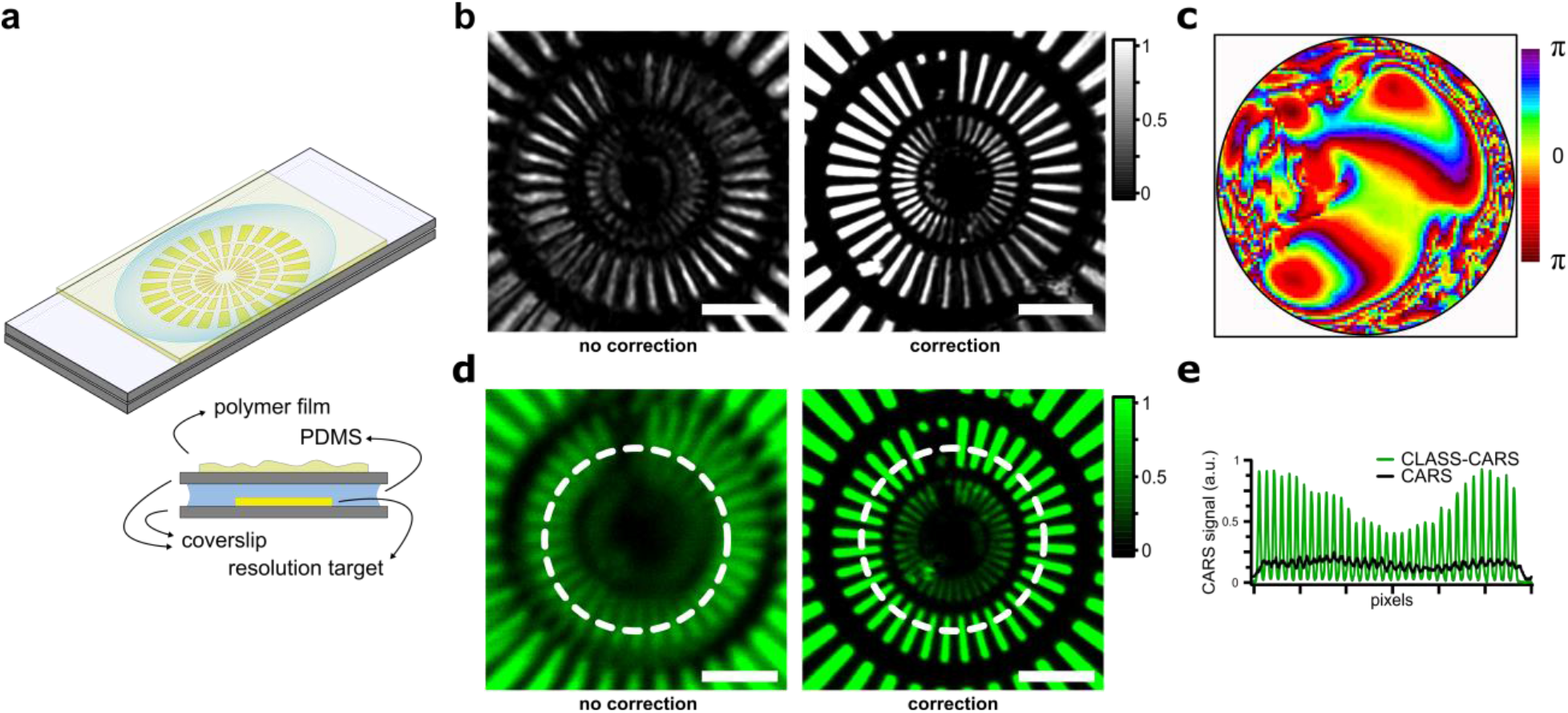
CARS imaging of an extended target under extremely severe aberration. **a,** Schematic presentation of the artificial target. A gold-deposit resolution target is immersed in PDMS polymer, and an irregular polymer film is placed on the top to introduce optical aberrations. **b**, Time-gated confocal reflectance images reconstructed by the reflection matrix before and after the computational aberration correction. Color bar: normalized by the maximum intensity of the corrected image. **c**, The phase map for aberrations in illumination and detection pupils retrieved by the CLASS algorithm, respectively. The radii of the maps correspond to a numerical aperture of 1.0. Color bar: phase in radians. The number of angular modes used for aberration correction in the pupil is about 6200. **d**, CARS and CLASS-CARS images, respectively. Intensity scales are set to the aberration-corrected images. **e**, Circular cut profiles along the white dotted lines in (d). Scale bars: 10 μm.

The complexity of the aberration maps in Fig. 3c is so high that the initial pump Strehl ratio is only 0.008, which means that the CARS signal generated at the focal plane can be lower by almost a factor of 10^5^ before the aberration correction. This observation explains why there is nearly no CARS signal from the fine structures in the central region before aberration correction. On the other hand, the enhancement of the signal intensity shown in Fig. 3e is much lower than the theoretically expected enhancement estimated by the pump Strehl ratio. This is partly because the CARS signal is generated throughout the volume of the thick PDMS layer covering the resolution target. For the 150-μm-thick PDMS sample, we observed that the CARS signal intensity is linearly proportional to the pump Strehl ratio with a slope of 0.87 (Fig. S5). This result indicates that the upper limit of CARS signal enhancement is lower than the simple consideration, ~10^2^ not 10^5^. Again, the CARS signals generated by the PDMS layer far from the focal plane are not enhanced as much by the aberration correction.^34^ Additional factors include the effect of the extended target whose dimensions are much larger than the diffraction-limited resolution and the correction efficacies of the SLM for two different wavelengths with complex aberration maps. It should be noted that the complexity of the aberration in this sample is well beyond the capacity of the previous sensorless AO systems.^22^ The aberration map in Fig. 3b consists of 6200 orthogonal angular modes, more than two orders of magnitude larger than the number of correction modes in the typical sensorless AO.

After thoroughly validating the performance of the AO-CRS microscope, we attempted to image neuronal structures in the mouse brain in situ. Previous CRS imaging studies conducted tissue imaging either for an excised tissue section or through a cranial window to ensure clean optical access.^10,35^ However, skull openings can cause brain inflammations and perturb tissue structures and physiological environment. To minimize these potential artifacts, we conduct mouse brain imaging through a thinned skull.^36^ Compact bone and spongy bone are removed from the skull of a 8-week-old mouse to reduce its thickness to 30-40 μm (Fig. 4a, see Methods for sample preparation). We recorded a time-gated reflection matrix for a field of view of 80 × 80 μm^2^ at a depth of 50 μm from the upper surface of the skull. In fact, even the thinned skull induces strong aberration that can devastate the CRS imaging. Furthermore, the extent of aberration varies from position to position because the thinned skull is located close to the depth of interest. To address these local aberrations, we divide the recorded view field (40 × 80 μm^2^) into 7×15 sub-regions with 15 μm overlaps with one another and identify aberrations for individual 20 × 20 μm^2^ sub-regions (Fig. 4b). Strehl ratio enhancement estimated by the aberration varies from 3 to 27, and the average Strehl ratio enhancement was 10 (Fig. S6). Figure 4c shows the time-gated confocal reflectance images reconstructed from the recorded reflection matrix before and after the computational aberration correction. Myelinated axons having high reflectance than the surrounding tissues are visible after the aberration correction.

**Fig. 4:**
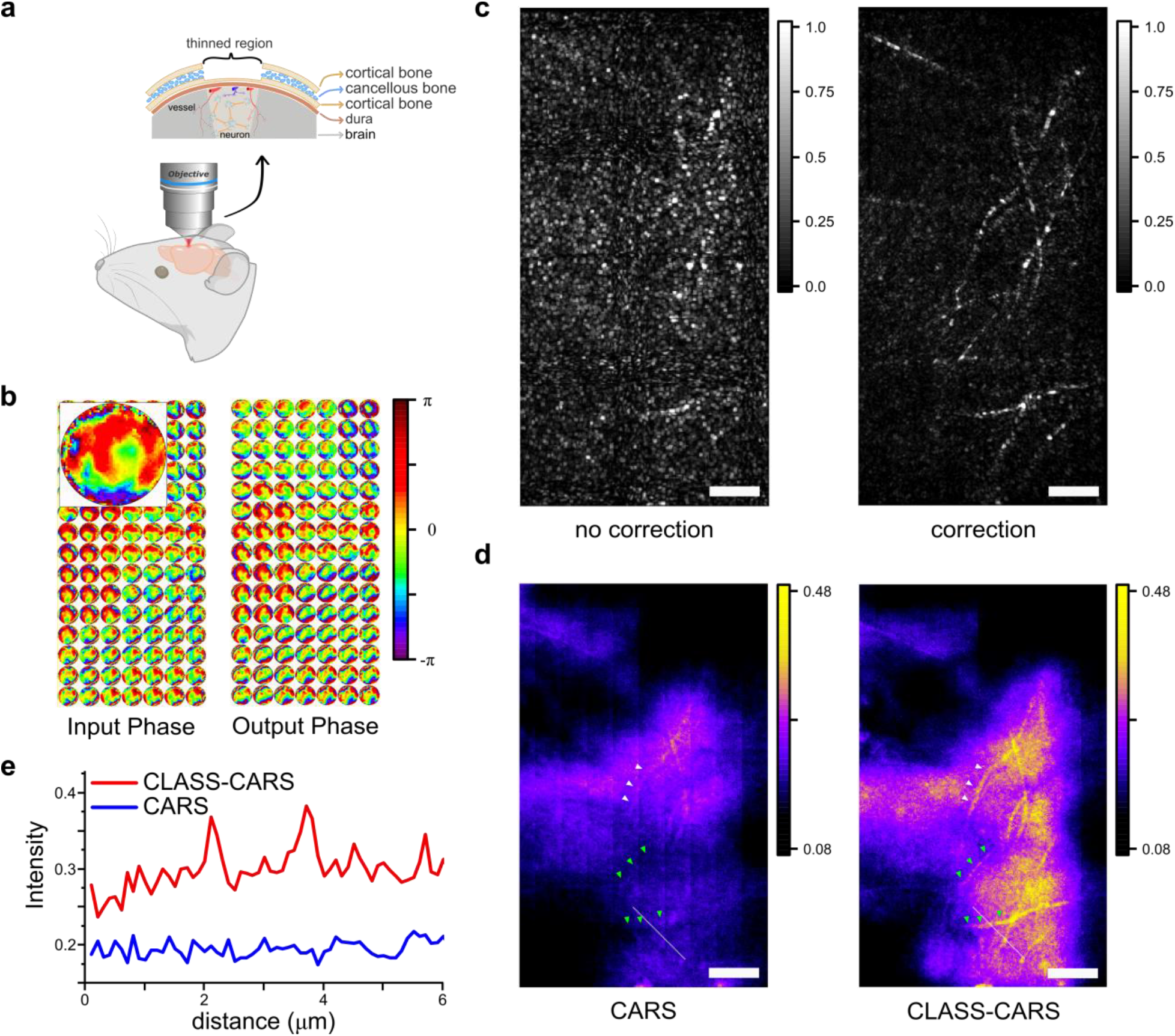
Coherent anti-Stokes Raman scattering images of mouse brain with the thinned skull. **a**, Schematic description of imaging configuration and the thinned-skull mouse sample. **b**, Input and output aberration maps measured by the CLASS system. Each circular map indicates the aberration map obtained for each sub-region. Zoom-in image: representative aberration map. Color bar: phase in radians. **c**, Time-gated reflectance images reconstructed by the reflection matrix before and after the aberration correction. **d**, CARS and CLASS-CARS images. The invisible myelin fibrils are made visible (green arrowheads) and the image contrast is significantly enhanced (white arrowheads). **e**, The line cut profiles along the dashed lines in (d). Scale bars indicate 10 μm in all the images.

With the aberration information at hand, we conduct CARS imaging of the lipid-rich substances by setting the energy difference of pump and Stokes beams matched with the CH vibration at 2850 cm^-1^. CARS images are obtained separately for individual sub-regions by displaying the aberration correction maps of the associated sub-regions on the SLM. A total of 105 CARS images are obtained in 20 × 20 μm^2^ sub-regions, which are merged to get a single full-field CARS image by selecting the maximum signal intensity at the overlapping area (Fig. 4d). With the aberration correction, many myelin fibrils initially invisible are made visible (green arrowheads in Fig. 4d), and the image contrast is significantly enhanced for those vaguely visible before the correction (white arrowheads in Fig. 4d). There is a resemblance between the reflectance image (Fig. 4c) and CARS image (Fig. 4d) because the dense lipid-rich structures are the primary source of contrast in both cases. However, the CARS image provides more chemical-specific information than reflectance imaging. Furthermore, the myelin in the reflectance image is subject to speckle noise due to multiple scattering signals with the same wavelength, but the CARS image uniformly visualizes myelin fibrils.

For the quantitative analysis, line cut profiles are plotted in Fig. 4e along the dashed lines in Fig. 4d. The two peaks associated with the myelin fibrils are clearly resolved with the contrast of 3.86 and 5.82 with respect to the standard deviation of the surrounding background. Since the initial SNR could not be measured due to the lack of resolving power, it was not possible to estimate the SNR enhancement by the aberration correction. The widths of the myelin segments were 203 nm and 284 nm, close to the diffraction limit accounting for the nonlinear excitation. It should be noted that the contrast of the image acquired with the thinned skull is comparable to that without any aberration (see CARS image of the tissue surface in Fig. S7). Even without any aberration and scattering, the intensity level of myelin is only 1.8 times higher than its surrounding region, which is similar to the deep tissue measurement obtained with the thinned skull (Fig. 4d). This low contrast suggests that signals from the lipids in the background tissues are comparable to those from the lipids in the myelin. Non-resonant signals in the CARS images are another factor contributing to the background signal. Essentially, CARS signals originate from the entire tissue volume as well as myelinated axons in the focal plane. Signal enhancement occurs mainly at the focal plane where the aberration correction is effective, and the contribution of background signal from other depths remains almost the same. Therefore, the apparent signal enhancement is lower than that estimated with the pump PSF, which is valid only for the isolated point particles (Fig. 2). In another perspective, the contrast of CARS images is intrinsically low in deep-tissue imaging such that even small PSF perturbation can wipe out the structures of interest. Therefore, PSF correction is critical to recovering the microscopic structures. In addition to the thinned skull imaging, we demonstrate the deep-tissue CARS imaging by correcting the aberration by the blood vessels (Fig. S8) and the tissue itself (Fig. S9).

To conclude, we developed label-free adaptive optical coherent Raman scattering microscopy for deep-tissue vibrational imaging. An advanced wavefront sensing AO was integrated with CRS microscopy to measure complex tissue aberrations from elastic backscattering, independent of CRS imaging, and correct the aberrations of the pump and Stokes pulses using a spatial light modulator. We demonstrated the coherent Raman imaging of the myelinated axons in the brain cortex through a thinned skull that obscures microstructures underneath. Many myelinated axons initially invisible were made visible with near-diffraction-limited spatial resolution, and the image contrast is on par with that observed without the skull. This study marks an important milestone in extending the penetration depth of CRS imaging. We anticipate further improvement of imaging depth and image quality by using longer excitation wavelengths and independent wavefront control of Stokes beam. Since the proposed method can be applied for multiple imaging modalities such as stimulated Raman scattering microscopy and transient absorption microscopy, it will allow the label-free mapping of lipids, proteins, myelin, and blood vessels deep within tissues.

## Supporting information

whole SI

## Methods

### Beam preparation

We prepared two pump and Stokes beams based on the optical backbone of coherent Raman scattering microscopy, reported previously.^37^ A femtosecond laser (Coherent, Chameleon Discovery) provided two laser sources of a tunable (680–1300 nm) and a fundamental (1040 nm) outputs with a repetition rate of 80 MHz. We set the energy difference between two outputs to the target molecular vibrations in the experiments. For controlling the intensities, two sets of polarizing optics (polarizing beam splitter, λ/2 waveplate) are installed in the beam paths, respectively. After the dichroic mirror for spatial overlap, the beam profiles of two beams are purified by focusing them to the pinhole (Thorlabs, 50 μm) and then collimated.

### CLASS (Reflection matrix acquisition and identification of sample-induced aberrations)

To obtain a reflection matrix, a sample is raster-scanned with a focused pump beam only over a field of illumination (FOI) with a sampling interval of half a wavelength of the pump laser. A set of complex-field images of the reflected pump waves for all scan points is measured by the CLASS microscope. The image size (field of detection, FOD) can be chosen appropriately depending on the PSF size of the reflected wave field, which limits the number of modes for aberration correction. For the thinned-skull sample used in the experiments, the FOD was chosen to be about 20×20 μm^2^. It takes 40 s to record a total of 34,596 images for an FOI of 75 × 75 μm^2^. The acquisition time can be reduced by reducing FOD size but at the cost of the available number of aberration correction modes. The use of a high-speed camera available in the market can reduce the acquisition time to less than a few seconds.

In image post-processing, a reflection matrix is constructed from the recorded images, and the CLASS algorithm^20^ is applied to the matrix to identify sample-induced aberrations. Practically, thick and complex biological samples induce position-dependent aberrations over a large FOI, which cannot be corrected simultaneously with a single pupil correction map. We estimate the size of the isoplanatic patch, over which the pupil aberration does not change significantly, in the thinned-skull sample is approximately 30 μm. It is noted that the isoplanatic patch for CARS imaging is much smaller than that for reflectance imaging since the CARS signal proportional to the square of the pump beam peak intensity. The isoplanatic patch size for CARS imaging of the thinned-skull sample, measured as the diffraction-limited background CARS signal in aberration-corrected CARS images. For a better correction of local aberrations, we divide the full FOI into 15 × 15 patches of each size 20 × 20 μm^2^ and find a local aberration map for each patch. The adjacent patches are overlapped by 7 μm for better CARS image stitching.

The CLASS algorithm on a GPU (NVIDIA RTX A6000) takes about 0.2 s to find the aberration map for each patch, and thus the total post-processing time for generating aberration correction maps is 45 s.

### CRS

In order to generate and collect the CRS signal from the sample, a water dipping objective with high NA and long working distance is utilised (Nikon Fluor, 60x w NA1.00 WD 2.0). After the high OD short-pass filter (Semrock FF01-637/7), the CARS signal is steered to the photomultiplier tube (Thorlabs, PMT2101), and it is sent to the data acquisition (DAQ) device (National Instrument, PCIe 6361). A home-built MATLAB code simultaneously controls the Galvano scanning mirrors, PMT, and sCMOS camera (pco edge 4.2 M – USB –HQ).

### Sample preparation for the artificial target

The spin-coated polystyrene beads (1 micron, Sigma-Aldrich) are placed in between two coverslips. On one side of the sandwich cell, a coat of nail varnish is added as an aberration layer. A gold-deposit resolution target is immersed in PDMS polymer in between coverslips. And polymer film is placed on the top to give an irregular optical aberration. The spontaneous Raman spectra for polystyrene (PS) and polydimethylsiloxane (PDMS) are presented with that of oleic acid (OA) as a reference (Fig. S9).

### Sample preparation for ex vivo mouse brain imaging through thinned skull

Eight-week-old C57BL/6 mice were deeply anesthetized with an intraperitoneal injection of ketamine and xylazine at 100 and 10 mg/kg, respectively, and decapitated. The scalps were completely removed. The heads were fixed with 4% paraformaldehyde at 4 °C for O/N and then washed with phosphate-buffered saline (PBS) in triplicate. After the heads were glued on 35-mm-diameter dishes, compact and spongy bones were ground. The 3-millimeter area in the center of the parietal bone was thinned using a 0.5-mm surgical carbide bur in a high-speed dental drill. Afterward, the bone was smoothed with a stone bur and a polishing bur to a thickness of ~35 μm. All animal experiments were approved by the Korea University Institutional Animal Care & Use Committee (KUIACUC-2019-0024).

## Acknowledgements

We thank Prof. S. C. Hong for helpful discussion and Dr. B. Cheon and Ms. S. Lim for their contributions to the initial setup of coherent Raman scattering microscopy.

## Author information

### Contributions

Conceptualization: MC, WC; Methodology: JML, SY, SK, YC; Investigation: JML, YS, SK, YC, WC, MC; Visualization: JML, YS, SK, YC; bio-sample preparation: JHH; Funding acquisition: MC, WC; Supervision: MC, WC; Writing – original draft: JML, YS, SK, YC; Writing – review & editing: JML, YS, SK, YC, WC, MC

### Corresponding author

Correspondence to Wonshik Choi and Minhaeng Cho.

## Ethics declarations

Competing interests

The authors declare no competing interests.

