## Supplementary material for "Adaptive Optical Coherent Raman Imaging of Axons through Mouse Cranial Bone": whole SI

### Supplementary Text

**Feasibility of stimulated Raman scattering (SRS) microscopy**: As depicted in Fig. S1, we located the detectors, photodiode (PD), and PMT for SRS and CARS, respectively, at the Fourier planes to collect the coherent signals efficiently. While the blue-shifted CARS signal is easily separated from the pump and Stokes beams by the dichroic mirror (DCM2), the SRS signal is distinguished by the beamsplitter since the SRS process induced the beam intensity alternation (stimulated Raman gain/loss). Since the incident laser beams experience the beam splitter twice before and after the sample in epi-type detection, we have to choose the beam splitting ratio for generating or detecting the signals efficiently. It should be noted that two polarizing beam splitters (PBS) are installed to combine the CLASS and CRS sections, which results in the limitation of incident beam intensities. Thus, in the present work, to prove the concept, we demonstrated the AO-CRS scheme mainly for measuring the CARS images even though the detection channel for SRS was installed. It will be possible to acquire the SRS images of objects in deep tissues with our AO-CRS set-up. Then, we anticipate that the AO-SRS microscopy would be of use to obtain CRS images without any non-resonant distortion when more intense incident beams are assured.

**Linearity of CARS signal in Bulk PDMS**:
Despite the fact that the CARS is a third-order nonlinear process, the intensity enhancement by eliminating aberrations shows approximately linear behavior in the volumetric sample. It can be rationalized by noting that the total powers are invariant under wavefront shaping. For the dummy sample of 150-μm-thick PDMS (Sylgard184, Dow Inc, 10 to 1 mix ratio), as the Strehl ratio increases, the normalized CARS intensity increases linearly with a slope of 0.87 (Fig. S5). Nevertheless, a more detailed quantitative analysis will be needed in the future since we utilized one SLM to correct the phase (spatial and temporal) mismatch between the pump and Stokes beams.

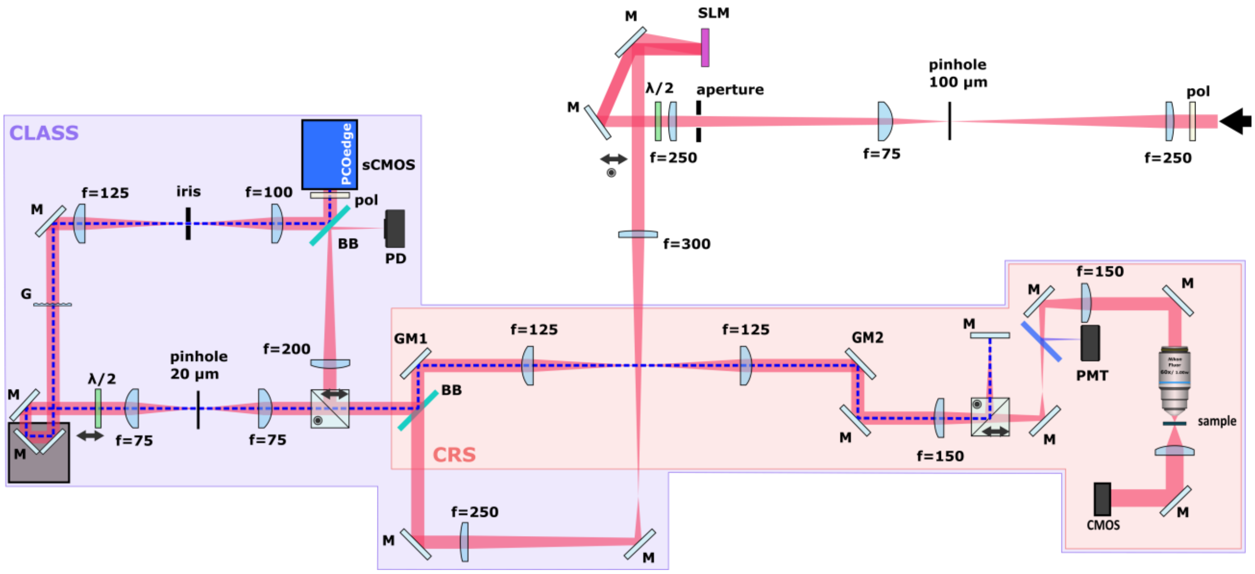


**Fig. S1. Detailed layout of the CLASS-CRS set-up.** pol: polariser, l/2: waveplate, M: mirror, SLM: spatial light modulator, BB: broadband beamsplitter, GM: Galvano mirror, G: grating, PD: photodiode. The focal lengths are indicated next to each lens. The photodiode is utilized to detect the SRS signal. CMOS camera or detector is located after passing through the sample to image the PSFs and conduct a transmission-type measurement.

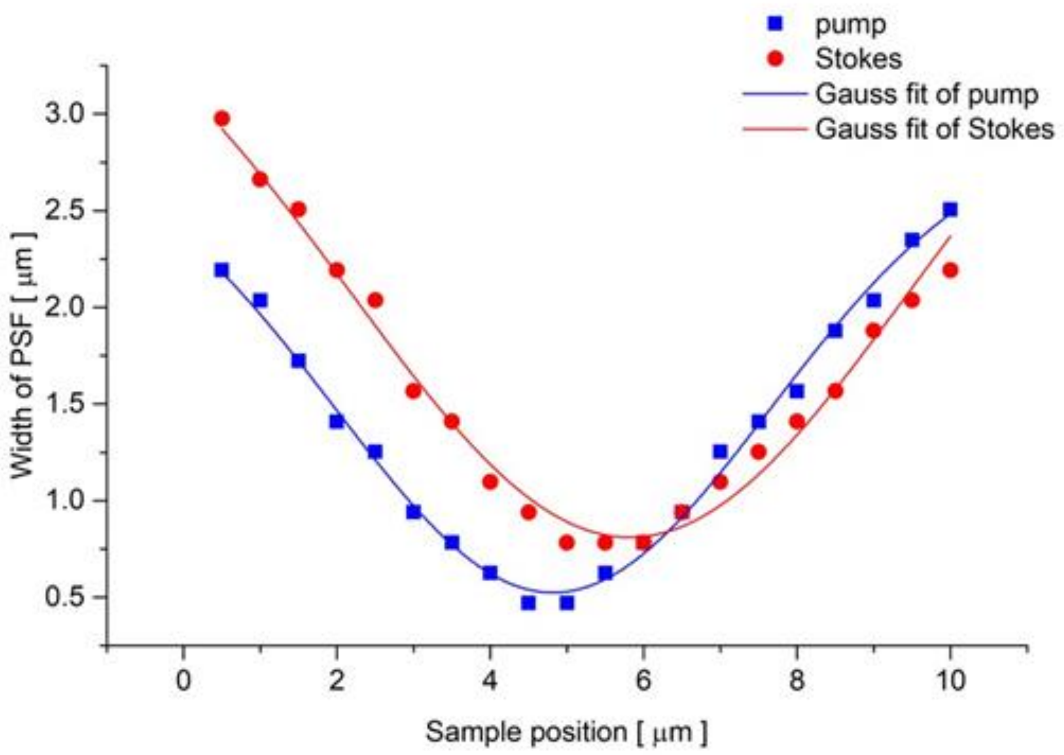


**Fig. S2. Z-position check for pump and Stokes beam.** The widths of the pump (blue) and Stokes (red) PSFs are measured with the knife-edge method at various z-position. The axial difference between the pump and Stokes focal points is about 1 μm.

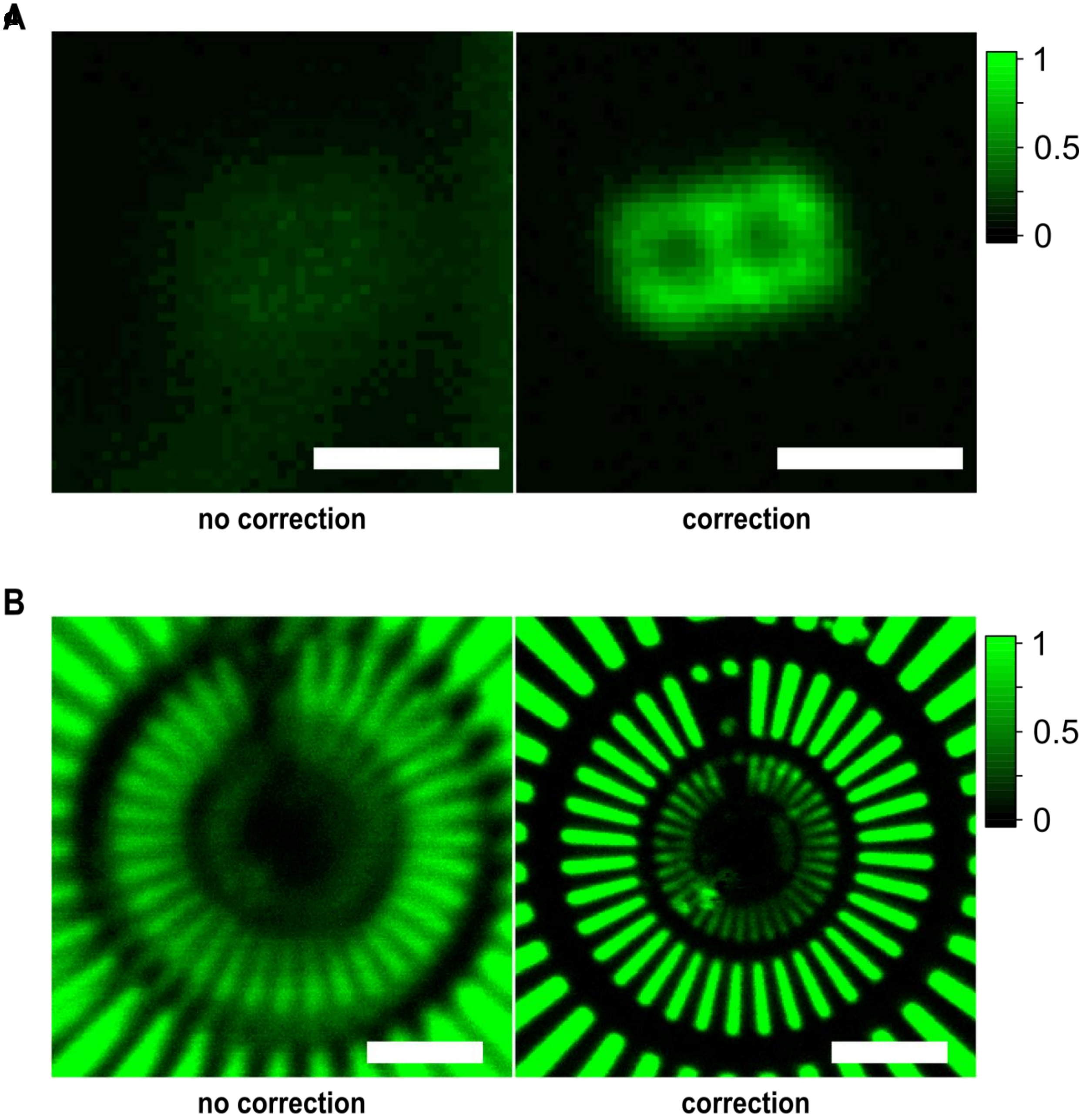


**Fig. S3. CARS imaging of an extended target under extremely severe aberration.** CARS images of (**A**) a resolution target indicator (figure-eight structure) and (**B**) resolution target (intensity scale magnified, x4) before (left) and after (right) correction. Scale bars: 2 and 10 µm for (A) and (B), respectively.

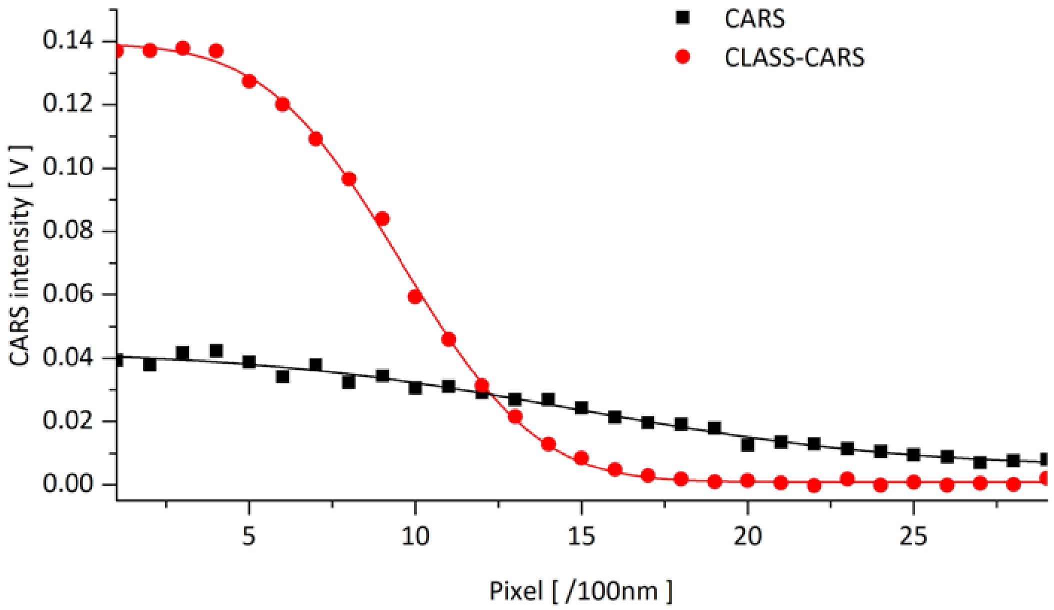


**Fig. S4. Line-cut profiles of CARS and CLASS-CARS images at the sharp edge of the resolution target.** PSF widths are estimated to be 1577 and 640 nm, respectively, supporting the enhancement of spatial resolution with the aberration correction.

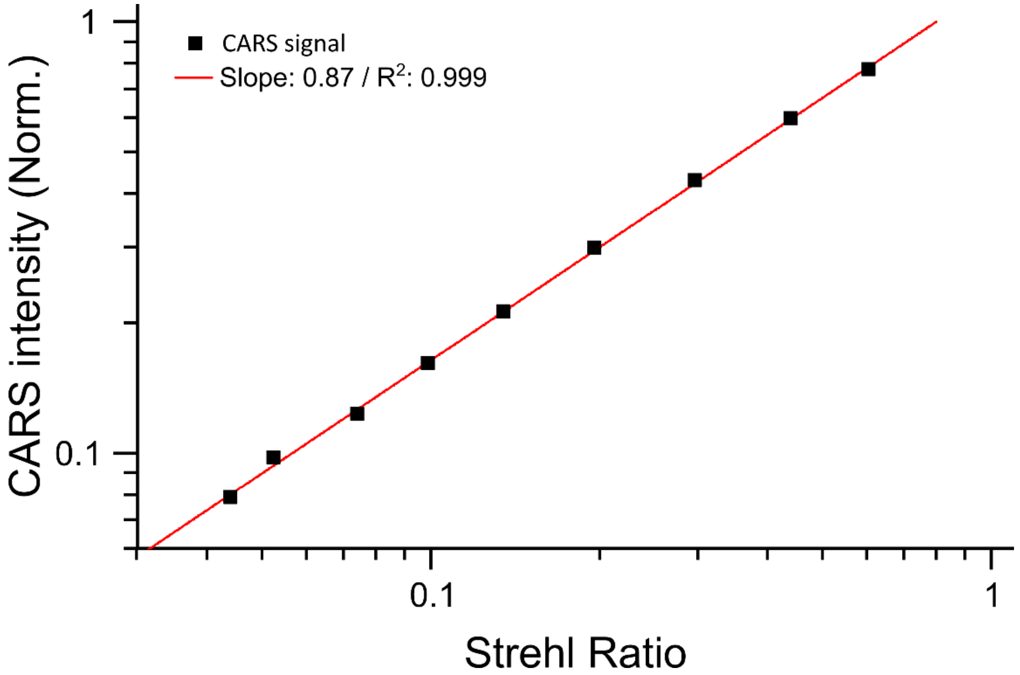

**Fig. S5. CARS intensity versus pump Strehl ratio.** The CARS signal intensity is plotted with respect to the pump Strehl ratio. The CARS signal is generated from the 150 μm-thickness PDMS layer.

A

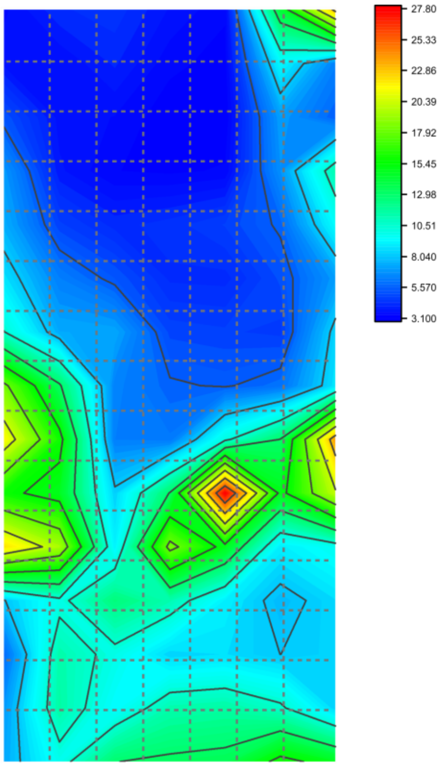


B

| | | | | | | |
|---|---|---|---|---|---|---|
| 3.53 | 4.29 | 4.49 | 3.81 | 3.40 | 14.17 | 23.37 |
| 3.77 | 3.60 | 4.14 | 3.43 | 3.23 | 9.27 | 6.38 |
| 5.26 | 3.74 | 3.58 | 3.16 | 3.12 | 7.06 | 5.58 |
| 6.93 | 3.90 | 3.36 | 3.29 | 3.39 | 6.79 | 11.42 |
| 7.43 | 4.90 | 4.24 | 4.46 | 4.85 | 5.70 | 9.45 |
| 8.67 | 6.34 | 5.33 | 4.28 | 4.24 | 5.14 | 7.01 |
| 10.55 | 7.31 | 7.25 | 4.99 | 4.88 | 4.73 | 8.39 |
| 18.28 | 13.19 | 6.44 | 5.67 | 5.49 | 5.97 | 8.73 |
| 21.51 | 16.00 | 6.31 | 6.12 | 10.22 | 13.02 | 24.00 |
| 15.85 | 14.71 | 8.27 | 13.06 | 27.76 | 14.91 | 20.12 |
| 22.28 | 19.53 | 9.08 | 18.48 | 14.83 | 8.94 | 9.79 |
| 8.00 | 10.10 | 12.57 | 11.33 | 8.99 | 7.63 | 8.51 |
| 5.76 | 11.42 | 10.04 | 8.34 | 8.56 | 8.04 | 8.40 |
| 6.92 | 11.58 | 9.31 | 11.38 | 11.62 | 10.75 | 8.34 |
| 7.29 | 9.89 | 12.95 | 13.24 | 13.46 | 15.94 | 14.77 |

**Fig. S6. The pump Strehl ratio enhancement estimated by the local aberration maps for mouse brain with a thinned skull.** (**A**) Strehl ratio enhancement map estimated with the aberration maps in Fig. 4B. (**B**) Enhancement factors at the corresponding subregions indicated by dashed boxes in (A).

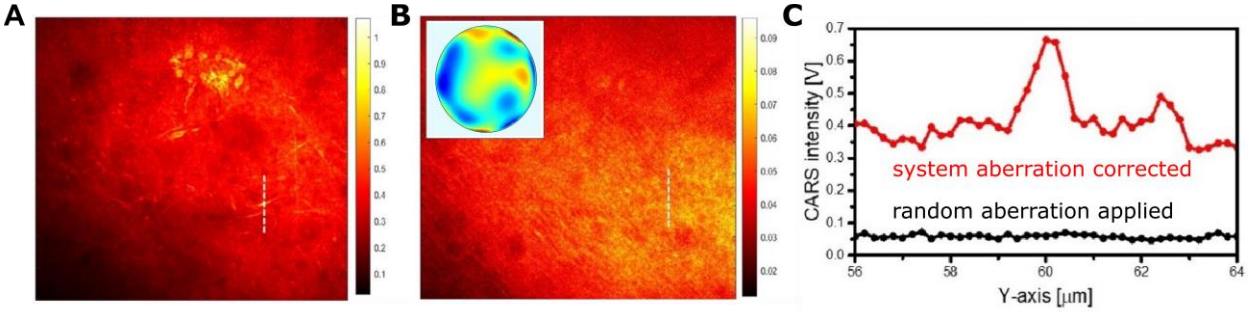


**Fig. S7. CARS images of myelin axons on the surface of the mouse brain tissue slice.** The grey matter region with sparse myelinated axons is imaged with CARS microscopy. (**A**) CARS image recorded after system aberration correction. (**B**) CARS aberration recorded after adding the artificial aberration map (inset) using an SLM. (**C**) Line profiles along the dashed lines in (A) and (B). Scale bar: 20 µm.

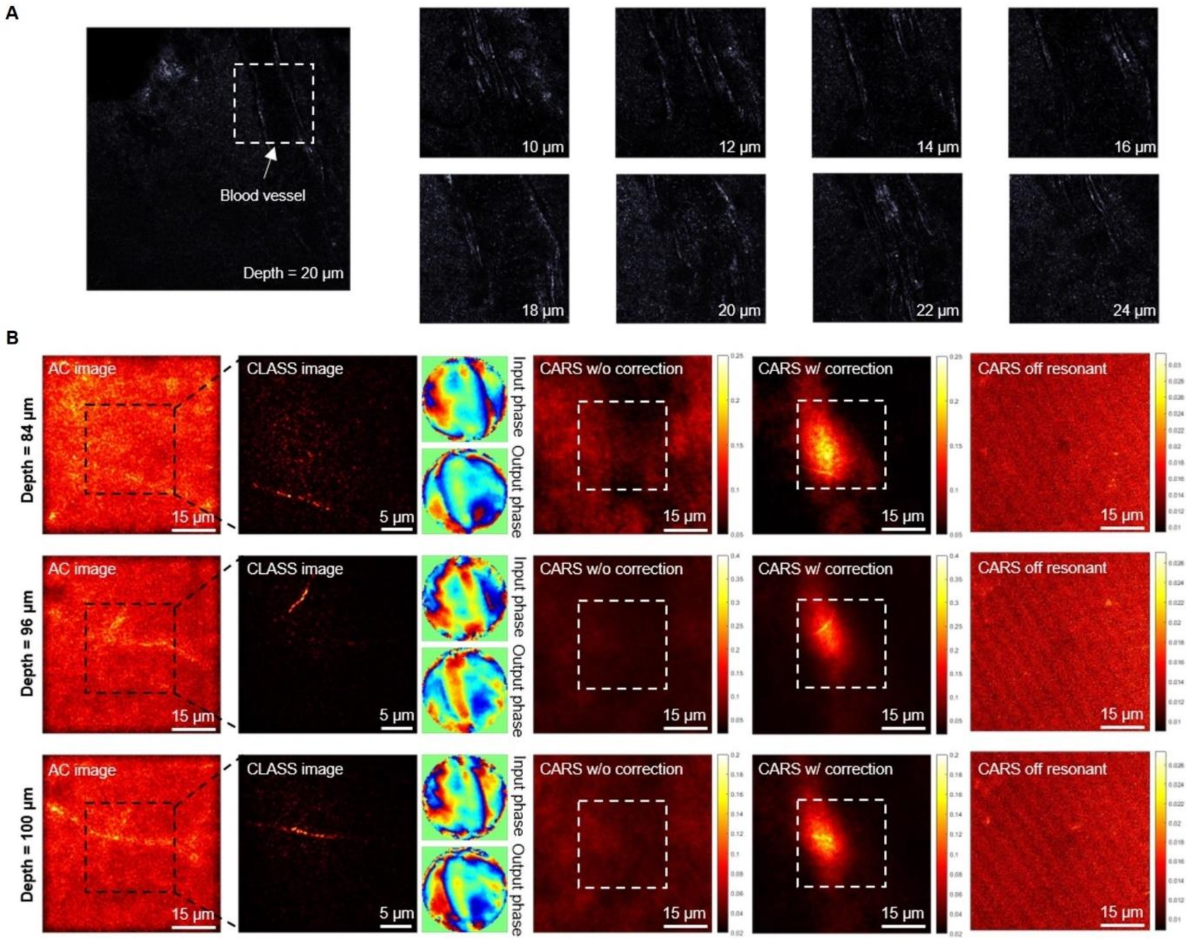


**Fig. S8. CARS images of mouse brain under a blood vessel.** (**A**) The reflectance images of a bare mouse brain in various imaging depths. The blood vessel is indicated by a white dotted box. (**B**) From the left, time-gated reflectance images reconstructed by the reflection matrix before and after the aberration correction, input and output aberration maps, CARS and CLASS-CARS images, and off-resonant CARS images, in three representative depths.

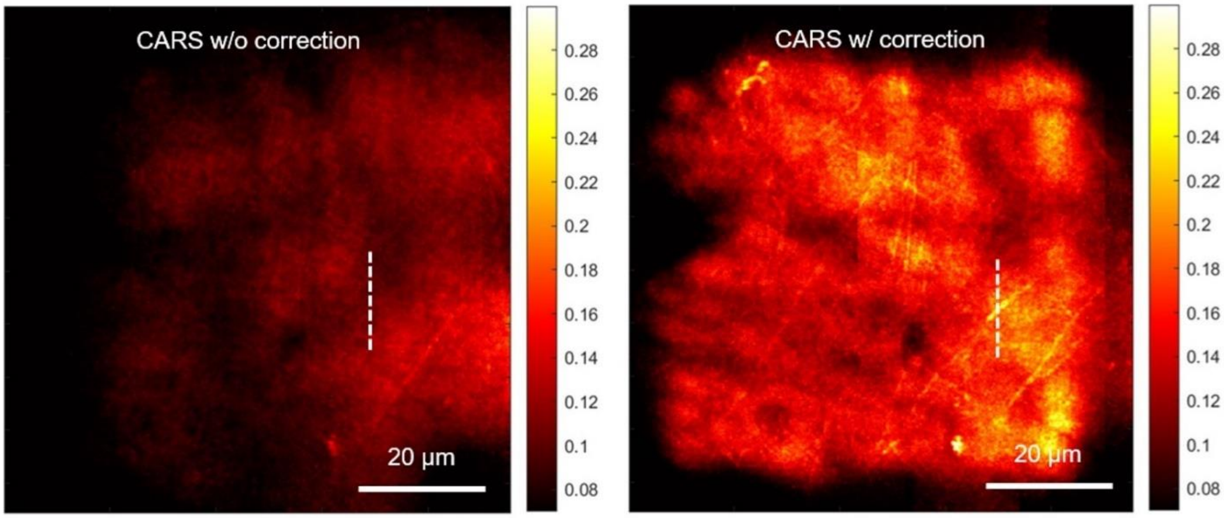


**Fig. S9. CLASS-CARS images of the whole mouse brain through a cranial window.** CARS (left) and CLASS-CARS (right) images. In the case of the CLASS-CARS image, local aberrations at 225 patches are corrected. Patch size is 20 x 20 μm, and each patch overlaps 15 μm.

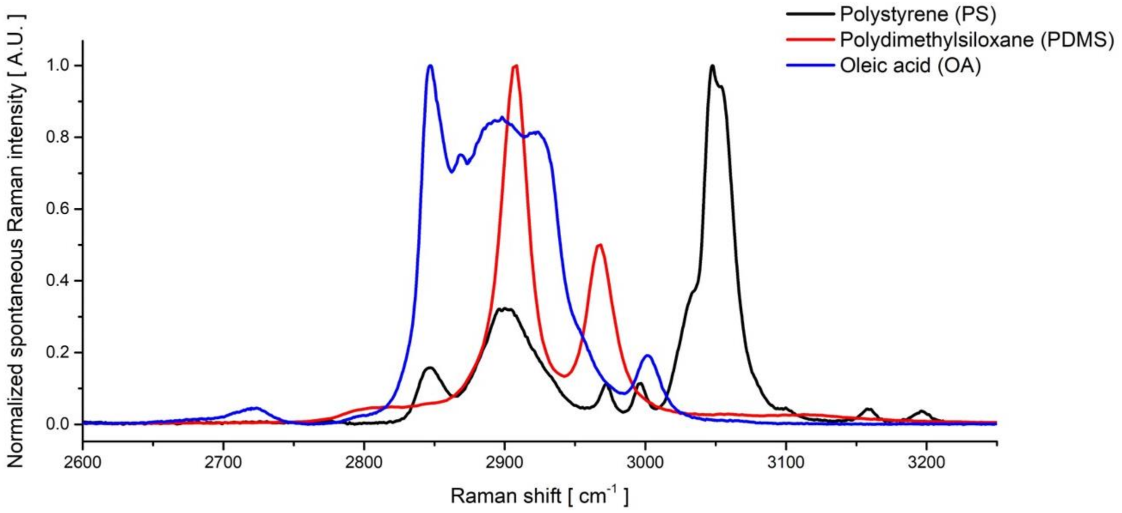


**Fig. S10. Spontaneous Raman spectra of three different chemicals.** Polystyrene (black), polydimethylsiloxane (red), and oleic acid (blue).
